# Human neurons stimulated with IFNγ present HLA class I-restricted autoantigens to cytotoxic CD8^+^ T cells

**DOI:** 10.64898/2026.05.22.727243

**Authors:** Benjamin DS Clarkson, Susanna Pucci, Ramila Barun Shrestha, Kiran K Mangalaparthi, Remya Raja, Marion Curtis, Akhilesh Pandey, Charles L Howe

**Author notes:** Correspondence to: Charles L Howe, PhD; Guggenheim 1542C; 200 First St SW, Rochester, MN 55905. The authors have declared no conflict of interest exists.

## Abstract

Interferon-γ (IFNγ) signaling is prominent in inflammatory CNS microenvironments across many neurological disorders, but the neuronal peptides presented on HLA class I under these conditions and their functional consequences for CD8^+^ T cells remain incompletely defined. Here we combine human iPSC-derived human neural aggregates (HNAs), HLA class I immunoprecipitation coupled to LC-MS/MS immunopeptidomics, and microfluidic co-culture assays to map IFNγ-induced neuronal antigen presentation and test antigen-specific cytotoxicity. IFNγ stimulation induced robust HLA class I expression in HNAs and enabled recovery of a canonical 8-12-mer class I ligandome enriched for 9-mers. Neuron-restricted expression of a synapsin-driven polyepitope cassette yielded presentation of defined neoantigen 9-mer peptides on donor HLA class I molecules and, in the presence of IFNγ, elicited activation of autologous antigen-specific CD8^+^ T cells and consequent antigen-dependent neurite injury. Across four donors, comparative immunopeptidomics identified large IFNγ-unique neural peptide repertoires distinct from matched fibroblasts and revealed a consistent enrichment of predicted high-affinity binders on HLA-B allotypes. Finally, β2-microglobulin deletion ablated peptide recovery, and neuron-restricted β2-microglobulin reconstitution enabled identification of neuron-derived peptides, including peptides derived from neurofilament light (NEFL) that were shared across donors and presented on multiple HLA allotypes. Together, these data provide an integrated platform for neuronal autoantigen discovery and functional validation and support a model in which IFNγ-driven neuronal HLA class I presentation creates an HLA-B-weighted epitope landscape that can be recognized by autoreactive cytotoxic CD8^+^ T cells.

## INTRODUCTION

Neuroinflammatory diseases of the central nervous system (CNS) are characterized by immune activation in close proximity to neurons and axons, raising the possibility of direct immune cell-neuron interactions that contribute to tissue injury [1]. In multiple sclerosis (MS), long-term disability correlates most strongly with gray matter pathology and neuroaxonal loss rather than lesion counts or relapse activity [2, 3]. Within active MS lesions, MHC class I-restricted CD8^+^ T cells predominate, outnumbering CD4^+^ T cells by an order of magnitude [4], and clonally expanded CD8^+^ populations persist across brain, CSF, and blood compartments, consistent with ongoing recognition of specific antigenic targets [4, 5]. Moreover, CD8^+^ T cell density within lesions correlates with active axonal injury [6], and lesion evolution and axonal injury track with clinical progression and disability accrual [7, 8]. Critically, CD8^+^ T cells are implicated in the pathogenesis, propagation, and/or progression of numerous CNS diseases, including Alzheimer’s disease, tauopathies, and ALS [9–14].

Cytotoxic CD8^+^ T cells mediate antigen-specific damage when target cells display peptides on human leukocyte antigen (HLA) class I molecules. Interferon-γ (IFNγ) is a key mediator of neuroinflammation that induces the antigen-processing machinery and HLA class I expression, potentially rendering neurons and axons more immunologically visible within inflamed microenvironments [1]. We previously showed that IFNγ drives retrograde induction of HLA class I molecules and antigen presentation machinery in human iPSC-derived neurons [15], and that CD8^+^ T cells recognizing neuron-restricted antigens can injure axons in a model of demyelination [16]. Together, these findings support the premise that inflammatory cues can enable neuronal and axonal antigen presentation sufficient for antigen-specific CD8^+^ effector injury, consistent with broader evidence that cytotoxic T lymphocytes participate in autoimmune and degenerative CNS diseases.

Despite evidence that inflammation renders neurons and axons susceptible to antigen-specific CD8^+^ T cell attack, the naturally presented peptide repertoire displayed by human neurons under inflammatory conditions remains poorly defined.Our prior work emphasized that neuron-specific autoantigens in MS remain unknown and may be difficult to detect using conventional approaches that prioritize dominant immune specificities [16]. In addition, inter-individual variation in HLA haplotypes raises the possibility that neuronal HLA class I ligandomes and the epitopes available for CD8^+^ T cell recognition vary substantially across individuals [1]. This point is underscored by evidence for tissue-specific HLA allotype expression/usage [17], locus-specific promoter differences in interferon-mediated induction of HLA class I genes [18], and genetic associations between specific HLA class I alleles and neuroinflammatory or neurodegenerative disease risk [19, 20]. Because HLA-A and HLA-B may also differ in the breadth and character of their peptide-binding repertoires [21, 22], direct measurement of neuronal peptide presentation across donors is likely to be essential for interpreting susceptibility and antigenic targeting in heterogeneous human disease.

Mass spectrometry-based immunopeptidomics provides a direct approach to identifying peptides naturally presented by HLA class I molecules and has been widely applied for antigen discovery [23–25]. Expression of antigen-processing genes or candidate proteins alone does not specify which peptides are processed and displayed on HLA class I molecules, reinforcing the need for direct ligandome measurement [23, 24]. Applying immunopeptidomics to neuronal systems requires scalable neural cultures, approaches to resolve neuron-specific contributions in mixed-lineage preparations, and functional assays that connect peptide discovery to antigen-specific cytotoxicity. Here, we establish an integrated platform combining human iPSC-derived neural aggregate (HNA) organoids with HLA class I immunoprecipitation and LC-MS/MS-based immunopeptidomics, alongside microfluidic co-culture assays using antigen-specific CD8^+^ T cells. We define IFNγ-induced HLA class I upregulation in HNAs, demonstrate neuron-restricted presentation of engineered neoepitopes sufficient to drive antigen- and IFNγ-dependent CD8^+^ T cell-mediated neurite injury, and map multi-donor IFNγ-induced neuroimmunopeptidomes to nominate candidate neuronal autoantigens for future evaluation as therapeutically relevant targets [1].

## RESULTS

### IFNγ stimulation induces expression of HLA-A/B/C by human iPSC-derived neural cells

Human neural aggregate organoids were prepared from healthy control and patient-derived induced pluripotent stem cells (iPSCs), as we previously described [15]. Briefly, neural stem cells (NSCs) induced by dual SMAD inhibition of iPSCs were differentiated using db-cAMP, BDNF, IGF1, GDNF, and B27 supplement. Analysis of cellular constituents in the organoids after 14-21 days of maturation using single nucleus RNAseq indicated enrichment for mature glutamatergic and GABAergic neurons, along with astrocytes, radial glia, and early and late neural progenitors (Figure 1A, 1B). Evaluation of βIII-tubulin, MAP2, and GFAP expression by immunolabeling and confocal fluorescence microscopy confirmed dense neuronal aggregates with extensive neurites situated within clusters of astrocytes (Figure 1C). We refer to these cultures as human neural aggregates (HNAs). Stimulation of HNAs with IFNγ (100 ng/mL) for 72 hours induced differential changes in expression of HLA class I genes relative to vehicle-treated HNAs, with a 2-fold increase in HLA-A (P=0.0004 by t-test; Hedges’ g = 2.6), a 3-fold increase in HLA-B (P=0.0005 by t-test; Hedges’ g = 2.4), and a 5-fold increase in HLA-C (P<0.0001 by t-test; Hedges’ g =6.0) (Figure 1D). This observation was confirmed by western blotting HNA lysates (Figure 1E) and by surface labeling for pan-HLA in unstimulated (Figure 1F) and IFNγ-stimulated HNAs (Figure 1G). We conclude that cultures of mature neural cells derived from human iPSCs upregulate expression of HLA class I molecules in response to stimulation with IFNγ, with particular enrichment at both the RNA and protein level for HLA-B and HLA-C expression.

**Figure 1.**
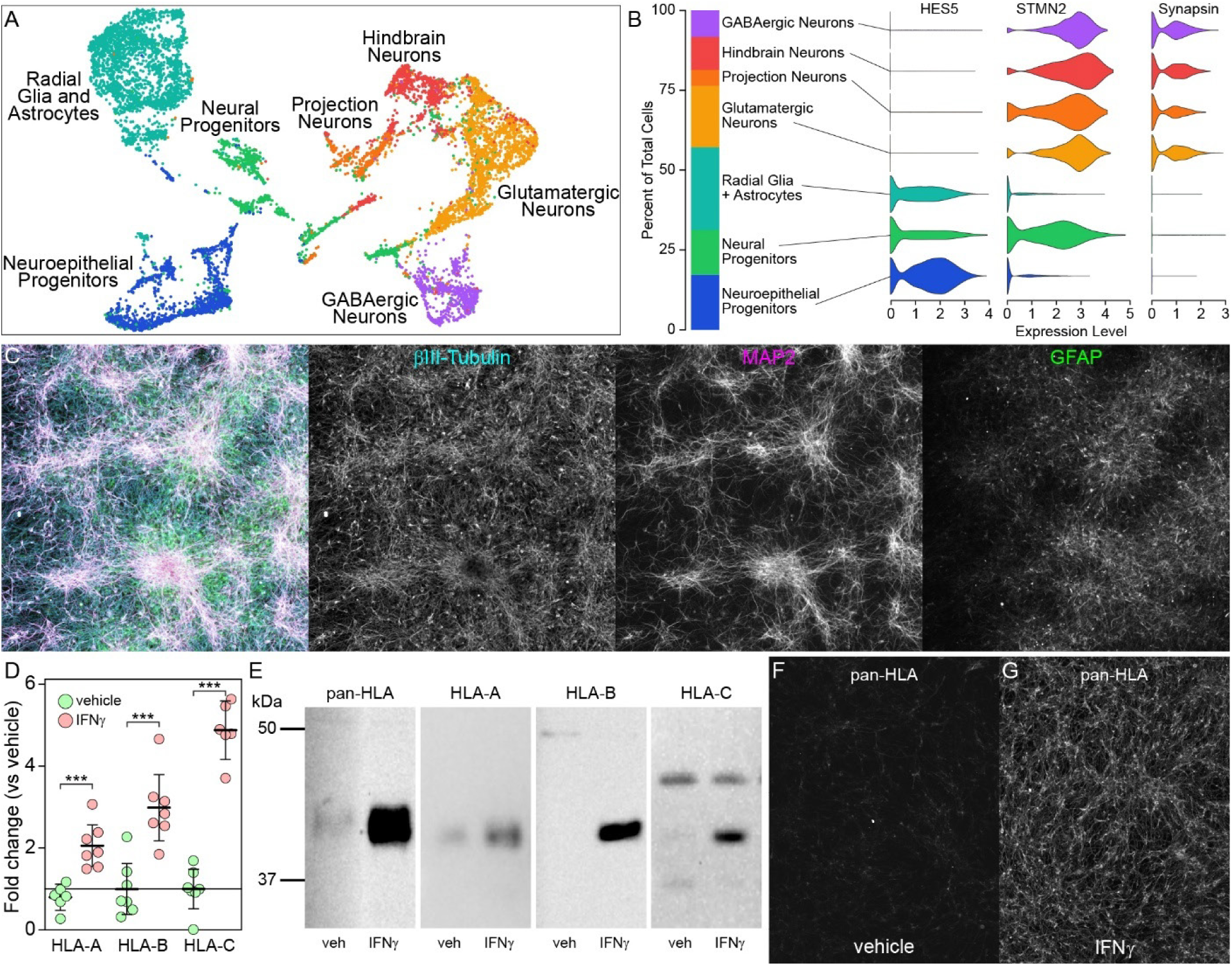
Characterization of the IFNγ-induced HLA class I response in human neural organoids. (**A**) UMAP visualization of single-cell RNA-seq from human neural aggregate (HNA) cultures, annotated into major neural lineage states including neuroepithelial progenitors, neural progenitors, radial glia/astrocytes, and multiple neuronal subtypes (glutamatergic, projection, hindbrain/spinal, and GABAergic neurons). (**B**) Cell-type composition (left) and expression distributions (right) for representative markers used to support annotation, including the progenitor marker HES5 and neuronal markers STMN2 and synapsin. (**C**) Representative immunofluorescence image of HNA cultures showing dense neuronal networks and mixed lineage composition, with single-channel views of βIII-tubulin (neuronal), MAP2 (neuronal dendritic), and GFAP (astrocytic) staining. (**D**) Quantification of IFNγ-induced surface HLA class I locus expression, shown as fold-change relative to vehicle across HLA-A, HLA-B, and HLA-C; each symbol represents an independent sample (n=6 or n=7) and error bars show mean and 95%CI; ***P<0.001 by pairwise t-test. (**E**) Immunoblot analysis of pan-HLA and locus-specific HLA-A, HLA-B, and HLA-C expression in vehicle- and IFNγ-treated HNAs. (**F-G**) Representative pan-HLA immunofluorescence images showing low basal class I staining in vehicle-treated cultures (F) and marked upregulation following IFNγ stimulation (G).

### Neuron-specific neoantigens are presented on HLA class I

To determine if expression of HLA class I molecules on neurons results in presentation of self-peptides, we engineered an adeno-associated viral (AAV) vector driving expression of defective ribosomal products (DRiPs) encoding immunodominant peptides from cytomegalovirus, Epstein-Barr virus, and influenza (CEF) downstream of the neuron-specific synapsin (SYN) promoter. Inoculation of HNAs with this AAV (AAV1.Syn.sig-CEF-DC-LAMP-T2A-eGFP.WPRE; hereafter designated AAV1.Syn.CEF-EGFP) (Figure 2A) or a control vector encoding EGFP downstream of the synapsin promoter without CEF (AAV1.Syn.EGFP) resulted in neuron-restricted expression of EGFP (Figure 2C, 2D). Exploiting this neuron-specific expression of neoantigens, we scaled up production of NSCs and differentiated HNAs to obtain >1.2 x10^8^ neural cells, stimulated with IFNγ for 72 h, and processed for immunoprecipitation of HLA class I:peptide complexes followed by LC-MS/MS-based analysis of HLA-bound peptides. This approach yielded 4364 peptides (restricted to 7-25 amino acids in length) from 2411 unique proteins. Notably, two CEF-derived 9-mer peptides were identified: VSDGGPNLY (CEF_374-382_) from the PB1 polymerase of influenza A and RPPIFIRRL (CEF_479-487_) from EBNA3 (Figure 2B). We analyzed these two peptides in NetMHCpan 4.2 for allele-specific binding [26] and defined eluted-ligand (EL) ranks <0.5 as “strong binders” and 0.5<EL<2.0 as “weak binders” [27].

**Figure 2.**
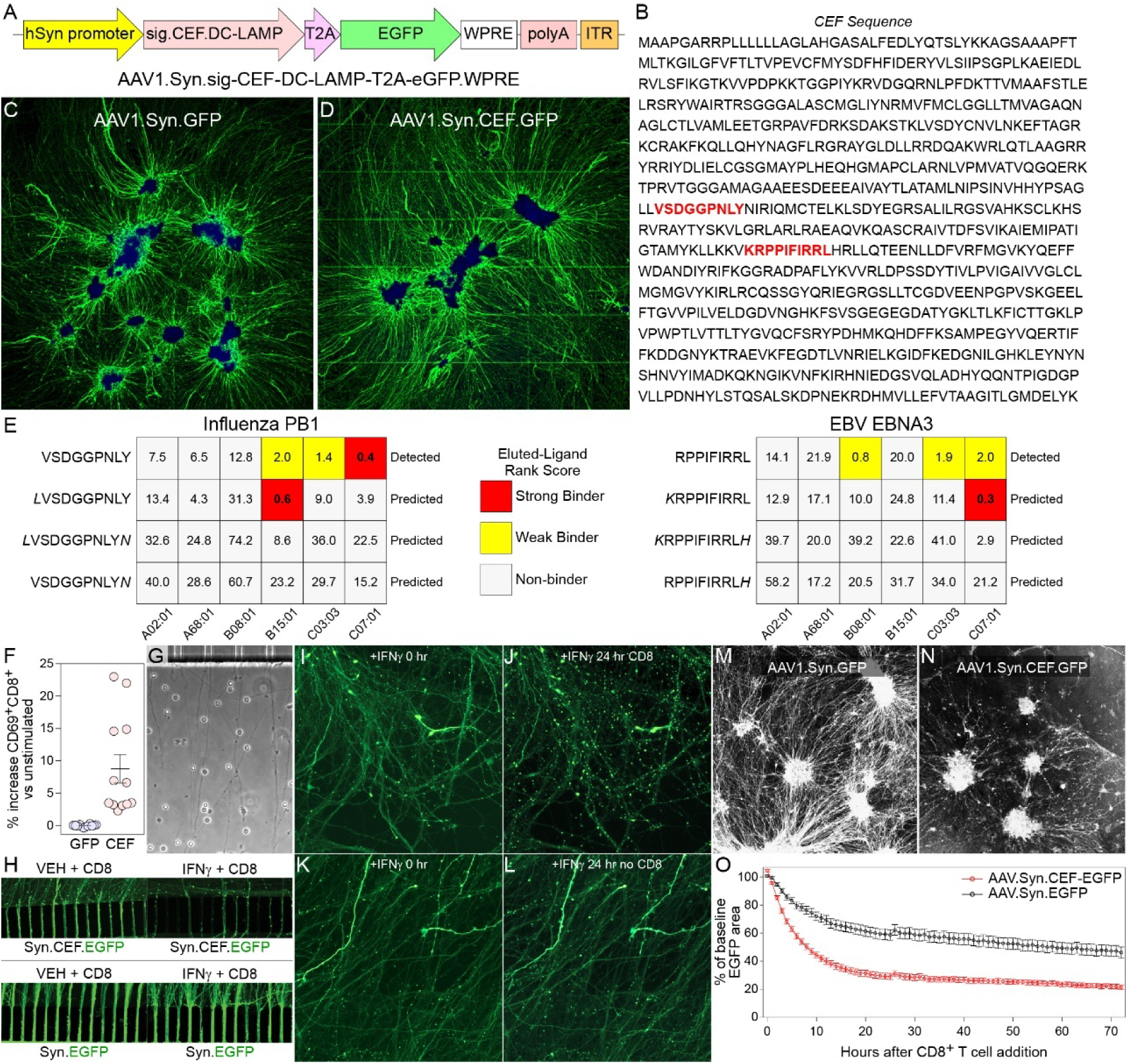
Neuron-restricted expression of defined neoepitopes drives antigen-specific CD8⁺ T cell recognition and consequent neuronal injury. (**A**) Schematic of the AAV expression cassette used to drive neuron-specific expression of a polyepitope antigen (CEF = CMV, EBV, inFluenza) under the human synapsin promoter (hSyn), coupled to EGFP via a self-cleaving T2A sequence (AAV1.Syn.sig-CEF-DC-LAMP-T2A-eGFP.WPRE). (**B**) Amino-acid sequence of the expressed antigen cassette with the embedded viral epitopes identified below (influenza PB1 and EBV EBNA3) shown in red. (**C**) Representative fluorescence images of control-(AAV1.Syn.GFP) and (**D**) CEF-transduced (AAV1.Syn.CEF.GFP) neural cultures demonstrating robust neuronal EGFP expression and neurite-rich morphology. (**E**) NetMHC-based eluted-ligand rank predictions for CEF epitopes (influenza PB1 and EBV EBNA3) identified by LC-MS/MS analysis of peptides bound to neural cell HLA class I following IFNγ stimulation; heatmaps summarize predicted binding class (strong/weak/non-binder) across donor HLA class I allotypes and detected peptides are indicated alongside predicted single amino acid extension variants consistent with antigen processing. (**F**) Quantification of antigen-specific CD8⁺ T cell activation following co-culture with control-(GFP) or CEF-transduced neurons, shown as the percent increase in CD69⁺CD8⁺ T cells relative to unstimulated controls; symbols indicate individual replicates and bars summarize mean and 95%CI. (**G**) Representative brightfield image demonstrating T cell interaction with neurites in the microfluidically isolated device co-culture condition. (**H**) Representative images showing antigen- and IFNγ-dependent neurite injury in CEF- or GFP-expressing neurons following addition of antigen-specific CD8⁺ T cells into the axon chamber of the microfluidic device. (**I-L**) Time- and condition-specific fluorescence images showing loss of neurite integrity only in the presence of IFNγ and antigen-specific CD8⁺ T cells. (**M-N**) Representative images of profound neurite loss following addition of antigen-specific CD8^+^ T cells into the cell body chamber only in the CEF-transduced cultures (**N**). (**O**) Longitudinal live-cell imaging quantification of GFP^+^ neurite loss, expressed as percent of baseline EGFP area over time following CD8⁺ T cell addition, comparing CEF-expressing (red) to control AAV-transduced (black) neurons.

Across the HLA class I haplotype of the healthy donor used for this experiment (HLA-A02:01, HLA-A68:01, HLA-B08:01, HLA-B15:01, HLA-C03:03, HLA-C07:01), the influenza peptide is predicted to bind strongly to HLA-C07:01 and weakly to HLA-C03:03 and HLA-B15:01 (Figure 2E). The EBV peptide is predicted to bind weakly to HLA-B08:01, HLA-C03:03, and HLA-C07:01 (Figure 2E). Because differentially trimmed versions of antigenic peptides may exist, we also analyze binding of ±1 terminal amino acid variants of the two detected peptides. This revealed that the N_term_+1 peptide KRPPIFIRRL from EBV (CEF_478-487_) is a strong binder to HLA-C07:01 and the N_term_+1 peptide LVSDGGPNLY from influenza (CEF_373-382_) is a strong binder to HLA-B15:01 (Figure 2E). We conclude that neuron-specific expression of CEF-derived peptides results in antigen processing and presentation on cell surface HLA class I.

### Neuron-specific neoantigens presented on HLA class I elicit CD8^+^ T cell-mediated injury

Given the broad immunity against EBV and influenza, we sought to determine whether CD8^+^ T cells from the autologous iPSC donor used to generate HNAs would recognize the peptides presented on HLA class I and injure the neurons. We transferred AAV1.Syn.CEF-EFGP or AAV1.Syn.EGFP transduced HNAs into a multicompartment microfluidic device that separates neuron cell bodies from distal axons, as previously described [15]. In parallel, we expanded T cells from the autologous donor by culturing peripheral blood mononuclear cells (PBMCs) in IL7 for 3 days and we generated dendritic cells (DCs) by culturing PBMCs in GMCSF+IL4 for the same period. Subsequently, DCs were incubated with a CEF peptide pool and co-cultured with the T cells for a further 3 days to drive expansion of CEF-specific T cells. During the same period the HNAs were stimulated with IFNγ (100 ng/mL) to drive HLA class I upregulation. CD8^+^ T cell activation by the CEF-presenting DCs was verified by flow cytometry (Figure 2F). CD8^+^ T cells were then isolated by negative magnetic selection and added to the axonal compartment of microfluidic devices containing autologous HNAs transduced with CEF-EGFP or EGFP only (Figure 2G). GFP^+^ axons were imaged repeatedly in the distal chamber, with evidence of extensive injury only in the IFNγ-stimulated cultures incubated with activated CD8^+^ T cells (Figure 2H-L). Critically, axon injury was strictly antigen-dependent: CD8^+^ T cells only substantially injured axons in HNAs transduced with the CEF construct and only in such HNAs treated with IFNγ (Figure 2H, 2J). HNAs only transduced with EGFP, even when treated with IFNγ, were resistant to CD8^+^ T cell-mediated injury (Figure 2H). Introduction of anti-CEF CD8^+^ T cells into the cell body chamber elicited even more robust axonal injury, but only in HNAs transduced with CEF-EGFP (Figure 2M, 2N). Serial live-cell imaging demonstrated progressive loss of neurites starting shortly after the addition of CD8^+^ T cells in the CEF-EGFP transduced HNAs but not in the EGFP transduced cells (Figure 2O). We conclude that human neurons are competent to present self-antigens on HLA class I following IFNγ stimulation, resulting in axon injury by antigen-specific CD8^+^ T cells.

### Neuron-specific autoantigens are presented on HLA class I

Using the same HLA immunoprecipitation assay and LC-MS/MS we characterized the immunopeptidome in unstimulated and IFNγ-stimulated HNAs and fibroblasts from 4 different donors. iPSC-derived NSC-derived human neural cells and donor fibroblasts were expanded to >1.2 x10^8^ cells, stimulated with IFNγ or vehicle for 72 h, and processed for immunoprecipitation of HLA class I:peptide complexes, as above. As expected, the majority of peptides isolated from the neural cells were between 8-12 amino acids in length, with a predominance of 9-mers (Figure 3A). Within each subject we compared conditions to identify peptides (8-12 residues) that were unique to the IFNγ-stimulated neural cells relative to IFNγ-stimulated fibroblasts and unstimulated HNAs and fibroblasts. Of 3016 total peptides in Donor 1 neural cells stimulated with IFNγ, 1904 were unique to the IFNγ-stimulated neurons and not detected in the unstimulated neural cells or the fibroblasts. Donor 2 had 5534 unique out of 7564 total; Donor 3 had 5953 unique out of 8324 total; and Donor 4 had 9556 unique out of 11305 total peptides. Reducing this analysis to the protein level, of 2199 total protein hits, Donor 1 had 732 that were unique to the IFNγ-stimulated neural cells. Of 4127 total proteins, Donor 2 had 1707 unique; of 4324 total proteins, Donor 3 had 1851 unique; and of 5287 total proteins, Donor 4 had 3939 unique hits (Figure 3B).

**Figure 3.**
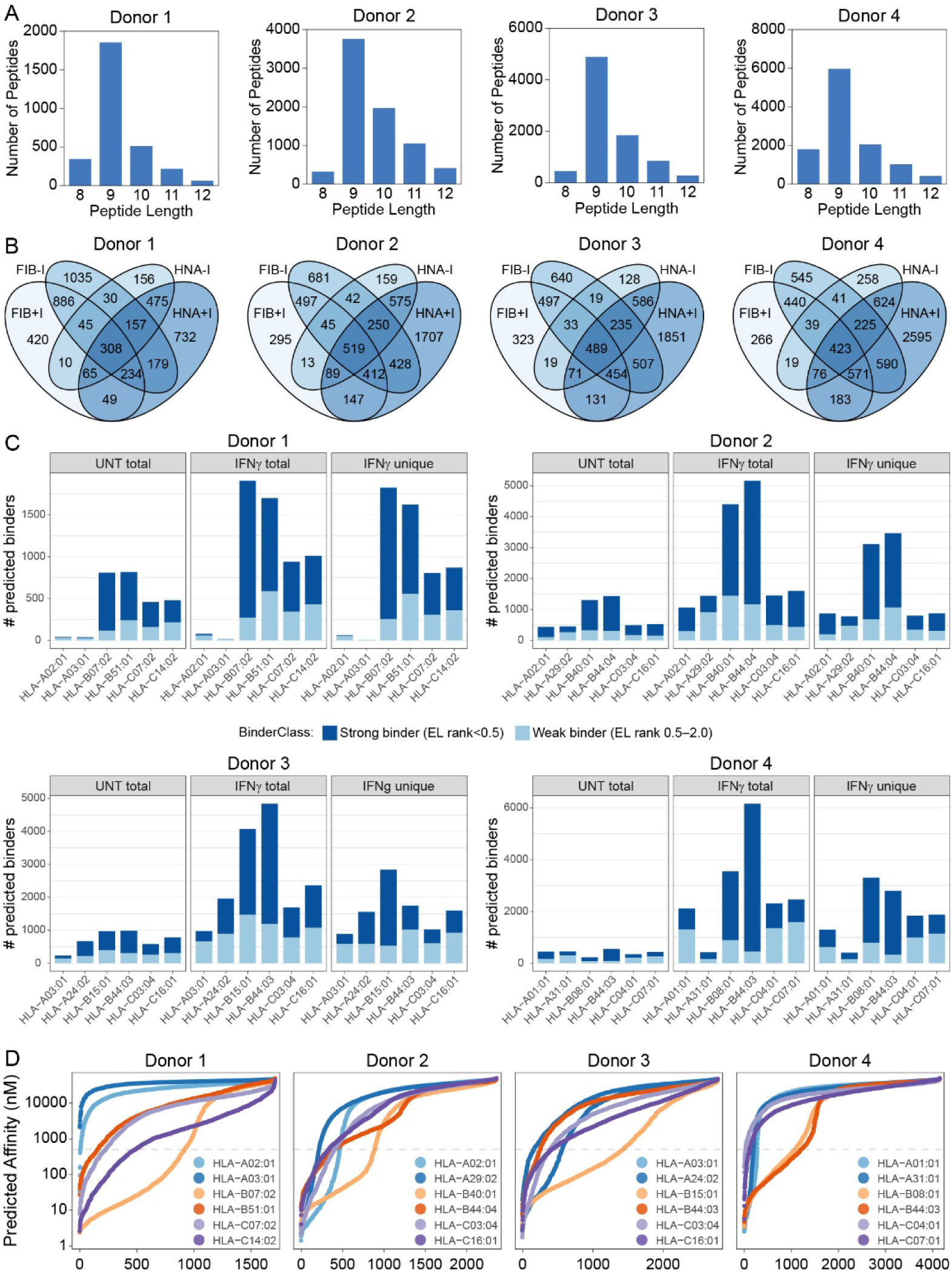
Multi-donor mapping of the IFNγ-induced neural immunopeptidome reveals reproducible HLA-B bias among predicted binders. (**A**) Length distribution of HLA class I-eluted peptides detected by LC-MS/MS from IFNγ-stimulated neural cultures across four donors, showing enrichment for 8-12-mers with a predominance of 9-mers. (**B**) Overlap analysis of antigenic protein repertoires across conditions within each donor, highlighting the subset of peptides unique to IFNγ-stimulated neural cultures relative to matched controls. FIB-I = unstimulated fibroblasts; FIB+I = IFNγ-stimulated fibroblasts; HNA-I = unstimulated neural organoids; HNA+I = IFNγ-stimulated neural organoids. (**C**) NetMHC-based binding classifications for donor-matched HLA class I allotypes, shown for untreated (UNT) total, IFNγ total, and IFNγ-unique peptide sets; stacked bars indicate predicted strong (dark blue) versus weak (light blue) binders by eluted-ligand rank thresholds, illustrating a consistent enrichment of predicted binders on HLA-B allotypes across donors. (**D**) Rank-ordered predicted affinity plots for IFNγ-unique 9-mers, stratified by donor HLA allotype; each curve represents peptides assigned to the best-predicted allotype, visualizing the high-affinity tail and the dominance of specific HLA-B allotypes in multiple donors; x-axis shows cumulative rank index of analyzed peptides; dashed line shows 500 nM cutoff.

NetMHCpan 4.2 was used to predict the affinity of each neural cell-derived peptide for the corresponding HLA class I allotypes present in each subject (Figure 3C). This analysis revealed that HLA-B-binding peptides outnumbered HLA-A and HLA-C binding peptides, both in the total immunopeptidome and in the peptidome unique to IFNγ-stimulated neural cells. Identification of strong binders (top 0.5%) and weak binders (top 2%) likewise indicated a preponderance of strong binding peptides on HLA-B for all 4 subjects. Analysis of the predicted affinity for the unique IFNγ-stimulated 9-mer peptidome for each subject using NetMHCstabpan 1.0 revealed that a single HLA-B allotype accounted for the majority of strong binding peptides in 3 of 4 donors (Donor 1, 3, 4), while the fourth donor showed strong binding to both HLA-B allotypes (Figure 3D).

Using CRISPR/Cas9 editing, β2-microglobulin (β2m) was deleted from Donor 4 iPSCs. According to inference of CRISPR edits (ICE) in the sequencing data from these cells we achieved >98% knock down of β2M. HLA immunoprecipitation and LC-MS/MS analysis of Donor4_β2MKO HNAs identified only 34 total peptides, of which 16 were 8-12-mers and 5 were 9-mers (compared to 6995 total, 6703 8-12-mers, and 4189 9-mers for the wildtype Donor 4 cells in this experiment). These peptides were likely derived from low-level contaminant carryover or non-specific binding. To refine the neuron-specific immunopeptidome within this framework, we reconstituted β2M expression in the Donor 4 β2MKO HNAs by transducing with AAV1.Syn.β2M. This led to the identification of 602 total peptides, 366 8-12-mers, and 141 9-mers derived from neurons. Of the 8-12-mers, 131 peptides were predicted as strong binders and 182 as weak binders using elute-ligand ranking. Comparing the predicted peptide binding affinity for the wildtype Donor 4 neural cells and the AAV1.Syn.β2M reconstituted β2MKO neural cells showed that HLA-B*08:01 dominated the high affinity pool in the wildtype (Figure 4A) while HLA-A*01:01 had the strongest representation in the reconstituted cells (Figure 4B). Because the reconstituted condition had ∼30-fold fewer total peptides scored for binding relative to the wildtype (141 vs 4189 9-mers used in the analysis), the distribution is likely undersampled, potentially skewing the HLA locus representation and missing rare high-affinity events. To facilitate a direct comparison between the wildtype and the reconstituted neural cell immunopeptidomes, we recast the by-allotype data as fraction of peptides with affinity <500 nM relative to the total number of peptides identified (Figure 4C). This normalization strategy revealed that the pattern of HLA-B enrichment in the wildtype was largely absent in the β2M reconstituted neurons and the high-affinity fraction was HLA-A weighted. To test whether the apparent loss of HLA-B predominance in Synβ2M could be explained by undersampling, we performed a depth-matched resampling analysis. Specifically, we repeatedly drew random sets of 141 WT 9-mer peptides (matching the Synβ2M 9-mer count), reassigned each peptide to its best-predicted HLA allotype (lowest predicted affinity), and quantified the resulting HLA-A/B/C distribution across 5000 iterations. Across all resamples, WT remained robustly HLA-B skewed and did not converge toward the Synβ2M locus distribution. While this suggests that the observed WT vs Synβ2M difference is unlikely to be an artifact of sampling depth, the approach does not exclude biological differences in antigen processing and loading or in peptide source pools between the WT condition and the neuron-restricted β2M rescue.

**Figure 4.**
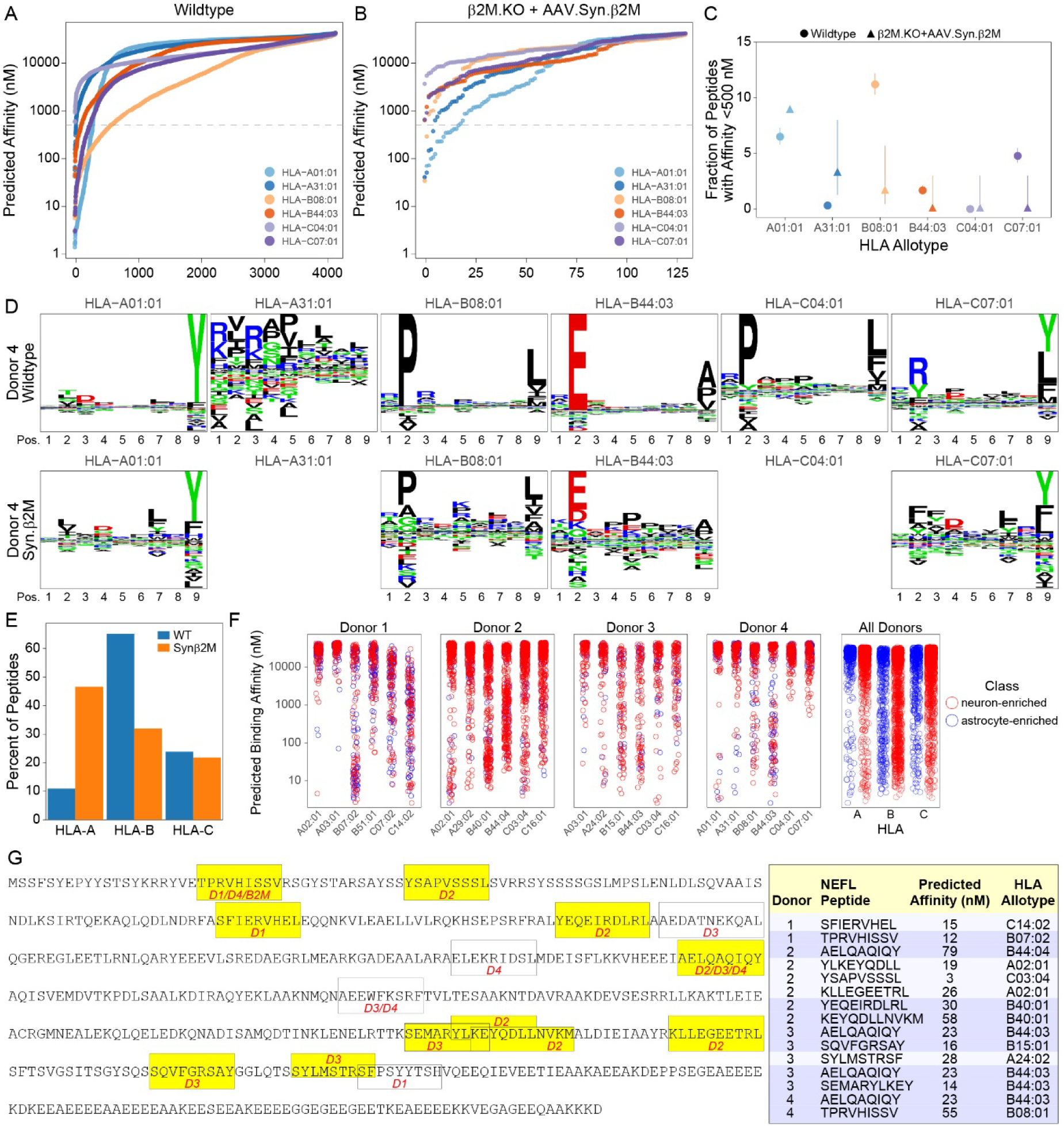
Neuron-restricted β2M rescue alters locus weighting and nominates several shared and unique neuronal autoantigens. (**A-B**) Rank-ordered predicted affinity plots for Donor 4 WT (**A**) versus the β2M knockout line from the same donor rescued with neuron-restricted β2M expression (β2M.KO + AAV.Syn.β2M) (**B**), showing the distribution of predicted affinities for HLA class I bound peptides across HLA class I allotypes; x-axis shows cumulative rank index of analyzed peptides; dashed line shows 500 nM cutoff. (**C**) Fraction of peptides below an affinity threshold (<500 nM) predicted to bind to each HLA allotype in WT versus the neuron-restricted β2M rescue, highlighting a shift in the high-affinity fraction across loci/allotypes. (**D**) Motif deconvolution of 9-mer repertoires shows inferred allele-restricted anchor preferences for each allotype between Donor 4 WT (top) and the neuron-restricted β2M rescue (bottom). (**E**) Summary of inferred locus composition (percent of peptides assigned to HLA-A, HLA-B, or HLA-C) comparing WT and neuron-restricted β2M rescue. (**F**) Predicted affinity scatter plots across donor-specific HLA allotypes for peptides derived from proteins classified by single-cell RNA-seq-informed enrichment (“neuron-enriched” versus “astrocyte-enriched”), shown for each donor and pooled across donors. (**G**) Example of a neuron-restricted antigen, neurofilament light chain (NEFL), shared by all four donors and presented in the neuron-specific β2M rescue condition; schematic protein map with detected peptides and donor sharing indicated; table inset summarizes representative NEFL-derived peptides with predicted affinity and best-assigned restricting HLA allotype.

To assess whether WT and Synβ2M peptidomes differed in allele-specific anchor preferences, consistent with differential HLA allotype binding, we deconvoluted the 9-mer repertoires using MHCMotifDecon 1.2 to infer likely HLA class I restrictions and to visualize anchor motifs (Figure 4D). We additionally summarized the inferred restriction fractions by locus (HLA-A vs HLA-B/C) and compared these estimates to NetMHC-based best-allele assignments (Figure 4E). To further test the hypothesis that neuron-derived peptides exhibit an HLA-A preference relative to glial cell-derived peptides, we used our single cell RNAseq data to assign a cell source classification (“neuron-enriched” vs “astrocyte-enriched”) to each peptide source protein identified by immunopeptidomics analysis of the four main donors. We then plotted the predicted affinity for each donor-specific HLA allotype (Figure 4F). Consistent with the IFNγ-unique immunopeptidome bias toward neuron-associated proteins, the affinity plots were dominated by neuron-enriched calls. However, across all 4 donors, we did not observe enrichment of HLA-A assignment for peptides derived from neuron-enriched proteins relative to astrocyte-enriched proteins (OR=1.08, Fisher’s P=0.51), and high affinity peptides from either cell classification were most frequently assigned to HLA-B allotypes (Figure 4F). Together with donor-to-donor variability in HLA-A representation for high-affinity assignments and the depth mismatch observed in the Synβ2M dataset, these results suggest that the WT neural immunopeptidome is robustly HLA-B skewed, while the apparent HLA-A-weighting in the Synβ2M condition is suggestive but should be interpreted cautiously (Figure 3, Figure 4).

Finally, despite donor-to-donor variability at the peptide level, during the allotype usage analysis we identified 51 neural proteins with peptides represented in the IFNγ-induced immunopeptidome of at least 2 of the 4 donors, with 6 present in three donors and 4 present in all four donors. None of the proteins were observed in the vehicle-treated immunopeptidomes under our filtering criteria. These shared neuronal proteins included synaptic and scaffolding factors (APP, CACNB4, CADM1, DTNBP1, GABRQ, GRIA4, KALRN, KIDINS220, MAP1A, MAP2, NLGN2, NETO2, NPTX2, SPTAN1, STON1, SYNE1, TAGLN3), axonal factors (KIF1A, MAP1B, NEFL, KCNQ2, RUFY3, AGTPBP1, INA, TUBB3), and guidance molecules (EPHB3, PLXNA3, PLXNA4, PLXNB2, DOCK7). The microtubule-associated kinase doublecortin-like kinase 1 (DCLK1), for example, was found in the peptidomes from all 4 donors, with predicted high-affinity binding to HLA-A*01:01 (6.3 nM), HLA-B*15:01 (29 nM), and HLA-C*03:04 (12 nM), amongst others. The neuron-restricted neurofilament light protein (NEFL) was highly represented in the immunopeptidomes of all 4 donors (Figure 4G), with predicted high-affinity binding on multiple HLA allotypes, including presentation on HLA-B molecules in all of the donors. Of particular note, NEFL was also identified in the immunopeptidome of the AAV1.Syn.β2M reconstituted β2MKO cells, with the peptide TPRVHISSV being found also in Donor 1. We conclude that IFNγ drives presentation of a broad set of neural antigens across donors and that WT HNA immunopeptidomes are consistently skewed toward HLA-B presentation. In one donor, neuron-restricted rescue of β2M expression shifts the high-affinity fraction toward HLA-A presentation, supporting the possibility that neuronal restriction can alter allotype usage, although additional depth-matched experiments will be required to fully separate sampling artifacts from biology.

## DISCUSSION

In this study we established an experimental framework for the analysis of neuronal HLA class I antigen presentation, the discovery of subject-specific neuroimmunopeptidomes, and the assessment of antigen-specific anti-neuronal CD8^+^ T cell-mediated injury. We showed that human iPSC-derived neural stem cell-derived aggregate organoids upregulate HLA class I expression in response to IFNγ stimulation (Figure 1) and that neurons in these organoids become competent to present peptides derived from neoantigens on class I following IFNγ stimulation (Figure 2). We also observed that this presentation has functional consequences, in that neoantigen-presenting neurons are targeted by antigen-specific autologous CD8^+^ T cells, resulting in neurite loss and neuronal injury (Figure 2). Using the same HLA immunoprecipitation approach, we characterized the immunopeptidome and predicted HLA allotype binding interaction in neural organoids from 4 different donors, revealing a pronounced skew toward HLA-B-mediated presentation of high affinity peptides across all subjects (Figure 3). We validated the neuron specificity of antigen presentation in our platform using deletion and neuron-specific reconstitution of β2-microglobulin and characterization of the IFNγ-stimulated neuroimmunopeptidome (Figure 4). Finally, we found that neurofilament light chain-derived peptides are ubiquitously presented on neuronal HLA class I across donors and across multiple HLA allotypes (Figure 4).

These findings extend our previous observation that IFNγ drives retrograde induction of HLA class I molecules and the antigen presentation machinery in human neurons [15] and that CD8^+^ T cells recognize neuron-restricted antigens in a mouse model of demyelination [16]. We characterized a large neural HLA class I ligandome that was comprised of canonical 8-12-mers and enriched for 9-mers, as expected for a class I immunopeptidome. We observed a robust IFNγ-induced repertoire that was unique from both the basal, unstimulated repertoire and from the immuopeptidome of autologous fibroblasts. These data are consistent with a model in which local neuroinflammation, as might be found in demyelinating lesions, around proteinopathic aggregates, and near other degenerative sites within the CNS [28] leads to not only upregulation of neural HLA class I molecules and the antigen loading machinery, but also to a remodeling of the peptide repertoire shaped by immunoproteasome activation [15, 16].

A prominent feature of our current findings was the bias toward HLA-B presentation across multiple donors with very different HLA haplotypes. Prediction-based allele assignment indicated that HLA-B-binding peptides outnumbered HLA-A- and HLA-C-binders in the total repertoire and particularly in the high-affinity subset (Figure 3). Locus-specific promoter differences in interferon-mediated induction of HLA class I genes [18] may explain HLA-B enrichment within the context of IFNγ-mediated inflammation, though it is notable that our neuron-specific β2-microglobulin reconstitution experiment suggested that HLA-A may also be relevant to neuronal peptide presentation. Previous reports have demonstrated that HLA-B alleles exhibit narrower, more restricted peptide-binding repertoires than HLA-A alleles [21], though others have demonstrated that HLA-B molecules present a larger number and greater diversity of unique self-peptides compared to HLA-A [22]. It is possible that these disparate findings are associated with differential affinity thresholds for peptide binding, with HLA-B representing a more curated, high-affinity repertoire relative to HLA-A. Our observations support this model, with analysis of binding motifs in the wildtype Donor 4 cells and the synapsin-specific β2-microglobulin reconstituted Donor 4 β2M KO cells showing preservation of the heavy-chain binding pocket selectivity (dominant anchors are largely conserved) but reduced binding stringency on HLA-B allotypes (marked by broader residue noise) in the neuron-specific results (Figure 4D).

In the pancreas, interferon signaling (IFNα) shifts the β-cell immunopeptidome from a resting HLA-A predominance to an HLA-B-restricted repertoire and patients with type 1 diabetes exhibit HLA-B hyperexpression relative to non-diabetics. Moreover, islet-infiltrating CD8^+^ T cells recognize granule peptides that are restricted to HLA-B [29]. Another study observed tissue-specific HLA class I allotype expression and usage, with some tissues showing HLA-B predominance relative to HLA-A [17]. The relevance of skewed HLA class I presentation to disease susceptibility and pathogenesis is unclear, since risk vs protection effects are highly context dependent (particularly with regard to HLA class II linkage). HLA-A*02 alleles, for example, are associated with markedly decreased risk of multiple sclerosis [19], while HLA-B*07:02 is associated with increased risk of Parkinson disease [20]. However, despite this complexity, the HLA-B bias carries important implications that extend beyond antigen discovery to therapeutic design principles. HLA-B allotypes exhibit differential dependence on the canonical peptide-loading machinery, with some allotypes showing resistance to TAP inhibition, resulting in a broader and more diverse peptide pool under stress or inflammatory conditions [30, 31]. In a CNS context where interferon signaling and proteostasis changes may reshape the processing and loading of self-peptides, such allotype-specific properties may contribute to genotype-dependent differences in the immunopeptidome sampled by autoreactive CD8^+^ T cells. This highlights the translational significance of understanding the genotype-, allotype-, and disease-specific elements of the neuroimmunopeptidome for the development of peptide-based tolerization, TCR-directed therapies, and antigen-specific monitoring assays to assess therapeutic efficacy [1].

Our β2-microglobulin perturbation experiments provide evidence of a neuron-specific peptidome that may have deep therapeutic relevance. We found that CRISPR/Cas9 deletion of β2M effectively produced a null HLA class I peptidome and created a powerful system into which we could reintroduce HLA class I expression only in neurons. The identification of peptides derived from neurofilament light chain protein across all four donors and in the β2M reconstituted neurons indicates that neuron-restricted autoantigens may be shared across disease states and genotypes. The recurrent presentation of NEFL-derived peptides is consistent with the clinical phenotypes of multiple sclerosis and other neurodegenerative disorders. The detection of neurofilament light chain in serum and CSF is widely interpreted as a marker of neuronal and axonal injury [32–35]. This factor is elevated early in inflammatory demyelinating disease states and elevation correlates with clinical measures of disease severity and progression rate across numerous CNS diseases. The observation of NEFL-derived peptides in the neuronal HLA class I ligandome suggests a mechanistic link between structural damage to neurons and immune visibility – namely, that proteins central to axonal integrity and function can become substrates for antigen processing and presentation on HLA class I in the context of injury and inflammation, triggering an autoimmune response. Moreover, presentation of NEFL-derived peptides on HLA class I is consistent with both the presence of anti-neurofilament antibodies [36, 37] and neurofilament-reactive T cells [38, 39] in multiple disease states. Our findings suggest that neurofilament light may be a key autoantigen that drives CD8^+^ T cell-mediated initiation, expansion, and/or progression of diseases ranging from multiple sclerosis to ALS and age-related neurodegeneration.

Several limitations constrain the interpretation of our findings. The wildtype neural organoids are comprised of mixed neural lineages, so the immunopeptidomes identified in this study must be interpreted as a composite neural HLA class I ligandome. This motivated our use of the neuron-restricted β2M rescue to identify a truly neuron-specific immunopeptidome, but this approach also was limited by a much smaller peptide repertoire. While this is likely reflective of true limits on the extent of neuron presentation, it does confound the precision in our locus usage analysis. Likewise, inference of allele restriction from prediction and motif deconvolution is certainly informative, but it is not equivalent to allotype-specific immunoprecipitation approaches and quantitative analysis of HLA-A/B/C abundance. Nonetheless, our platform and the identification of a reference neuroimmunopeptidome provides a path for future work aimed at establishing the pathogenic significance of these candidate neuronal autoantigens. Further deep immunopeptidomic discovery using our platform can be used to nominate population-level and individual patient-level proteins for testing as CD8^+^ T cell targets in our co-culture system. In the context of MS, where compartmentalized, clonally expanded CD8⁺ T cells are posited to contribute to progressive axonal damage [1, 16], linking a defined neuronal peptide-HLA complex to CD8⁺ T cell effector function would represent a decisive step toward establishing causal antigen specificity and would identify a new therapeutic target. Extending this same logic to other neurodegenerative diseases characterized by neuronal and axonal stress and neuroinflammation may enable the identification of novel pathogenic CD8+ T cell-mediated mechanisms that will create new opportunities for therapeutic intervention.

In summary, this study defines an integrated strategy for neuronal autoantigen discovery and functional validation. The wildtype IFNγ-induced neuroimmunopeptidome is robustly HLA-B skewed across several donors, and β2M knockout plus synapsin-specific β2M reconstitution provides evidence for neuron-specific antigen presentation, including the recovery of NEFL-derived peptides as a potentially “universal” neuron-specific autoantigen. Within the context of existing paradigms for the initiation and progression of CNS autoimmunity, our results support a model in which interferon-driven neuronal autoantigen presentation creates an HLA-B-weighted epitope landscape for autoreactive CD8^+^ T cells that may be amenable to targeted therapeutic interventions ranging from immunotolerization to anti-TCR CAR-T treatments.

## MATERIALS AND METHODS

### Human specimens

All human samples were obtained from individuals providing written informed consent following protocols approved by the Mayo Clinic Institutional Review Board (IRB 17-004547; IRB 13-007298), the Mayo Clinic Biospecimens Committee, and the Mayo Clinic Stem Cell Research Oversight Subcommittee. The iPSC-derived neural organoids used for snRNAseq analysis were from a healthy donor (54-year-old white, non-Hispanic female). The iPSC-derived neurons and autologous CD8^+^ T cells used for the CEF experiments were from two healthy donors (31-year-old white, non-Hispanic female; 34-year-old white, non-Hispanic male). The iPSC lines used for deep immunopeptidomics were: donor 1 = 73-year-old white, non-Hispanic female; donor 2 = 74-year-old white, non-Hispanic male; donor 3 = 56-year-old white, non-Hispanic female; donor 4 = 34-year-old white, non-Hispanic male.

### iPSC cultures and neural differentiation

Induced pluripotent stem cell clones were generated using Sendai virus reprogramming (CytoTune 2.0, Thermo Fisher) of adult dermal fibroblasts derived from skin punch biopsies. Trilineage potential, pluripotency, and sterility were assessed by the Mayo Clinic Biotrust and the Mayo Clinic Stem Cell and Organoid Core Facility. iPSCs were differentiated into neural stem cells using dual SMAD inhibition and STEMdiff Neural Induction Medium (Stem Cell Technologies). Following neural induction, NSCs were cultured in NSC expansion media consisting of KO DMEM/F-12, 1% Glutamax, 2% StemPro Neural Supplement (Thermo Fisher), 1% Anti-Anti (Thermo Fisher) and 20 ng/mL FGFb (PreproTech) and EGF (Invitrogen). iPSCs and NSCs were supplemented with 5 µM Rock Inhibitor (Y27632, Stemcell Technologies) for the first 24 h after thawing. Human neural aggregate organoids were differentiated in media composed of KO DMEM/F-12, 2% B27 supplement (Thermo Fisher), 1% Glutamax, 100 U/mL penicillin, 100 µg/mL streptomycin, 100 µM dibutyryl cyclic-adenosine monophosphate (cAMP; Sigma), 5 µg/mL plasmocin (Invivogen), 1 mM sodium pyruvate, and 10 ng/mL each of BDNF (PreproTech), IGF1 (Corning) and GDNF (PeproTech) for at least 2 weeks, as previously described [15].

### T cell coculture

Autologous cryopreserved human PBMCs prepared as previously described [40] were thawed, washed, and allowed to rest at 37°C in RPMI containing 10% heat-inactivated human serum. For CEF stimulation monocytes were isolated using EasySep Human Monocyte Isolation Kits (StemCell Technologies) and differentiated into dendritic cells over 72 hours with GMCSF (50 ng/mL) and IL4 (20 ng/mL) as previously described [40]. CD3^+^ T cells were purified from PBMCs using EasySep Release Human CD3 Positive Selection Kits (StemCell Technologies) and rested for 72 hours in 10 ng/mL IL7. DCs were pulsed with CEF peptide pools (10 µg/mL) and cocultured with CD3^+^ T cells for 72 hours. T cell activation was assessed by flow cytometry (Attune Nxt) using antibodies against CD3 (clone HIT3a), CD4 (clone OKT4), CD8 (clone HIT8a), and CD69 (clone FN50). CEF-stimulated CD8^+^ T cells were purified (EasySep Human CD8^+^ Isolation Kit) and added to autologous human neural aggregate organoids (5×10^5^ T cells per well of a 96 well plate). Automated live cell imaging and analysis was performed on an IncuCyte SX5 Green/Orange/NIR or IncuCyte SX3 Green/Red (Sartorius) with the neurite analysis module (Neurotrack, Sartorius). The coculture starting timepoint was used in every experiment as a reference value for signal normalization.

### Single nucleus RNA sequencing

Human neural aggregate organoids were processed using the Chromium Single Cell 3′ Gene Expression platform (10x Genomics) according to the manufacturer’s instructions. Dounce homogenization was performed on ice in nuclei isolation buffer containing Tris-HCl (pH 7.4), NaCl, MgCl₂, Nonidet P-40, DTT, and RNase inhibitor. Homogenate was filtered through a 40 μm cell strainer to remove debris and centrifuged at 500 × g for 5 minutes at 4°C to pellet the nuclei. Nuclei were resuspended in PBS supplemented with 1% BSA and RNase inhibitor, stained with trypan blue, counted using an automated cell counter, and adjusted to a final concentration of approximately 700-1,200 nuclei/μL. Single-nucleus suspensions were loaded onto a Chromium Next GEM Chip targeting recovery of 5000-10000 nuclei per sample. Nuclei were partitioned into Gel Bead-in-Emulsions (GEMs) using the Chromium Controller. Within each GEM, polyadenylated RNA transcripts were reverse transcribed using barcoded oligonucleotides containing a cell-specific barcode and unique molecular identifier (UMI). Following reverse transcription, GEMs were broken, and barcoded cDNA was recovered and amplified by PCR. Amplified cDNA was purified using SPRIselect magnetic beads.

Sequencing libraries were generated by enzymatic fragmentation, end repair, A-tailing, adaptor ligation, and sample index PCR according to the manufacturer’s protocol. Library quality and fragment size distribution were assessed using a TapeStation, and libraries were quantified by fluorometric and qPCR-based methods. Libraries were sequenced on an Illumina platform using paired-end sequencing with a 28-bp read for cell barcode and UMI capture and a 90-100-bp read for transcript sequence. Sequencing depth targeted approximately 30000-50000 read pairs per nucleus.

Raw base call files were demultiplexed using bcl2fastq and processed with Cell Ranger (10x Genomics). Reads were aligned to the appropriate reference genome using a pre-mRNA annotation to capture both intronic and exonic reads, enabling quantification of nuclear transcripts. Gene-barcode matrices were generated using default parameters. Downstream analyses were performed in Seurat in R. Low-quality nuclei were excluded based on gene complexity (>200 and <6000 detected genes) and mitochondrial transcript content (<15% mitochondrial reads). Doublets were removed using DoubletFinder. Data were log-normalized and scaled, 2000 highly variable genes were identified using the variance-stabilizing method, and principal component (PC) analysis was performed using 50 PCs. Dimensionality reduction and clustering were performed in Seurat using the first 20 PCs followed by shared nearest neighbor graph construction (FindNeighbors) and Leiden clustering with resolution set to 0.4. Uniform Manifold Approximation and Projection (UMAP) was used for two-dimensional visualization of transcriptional relationships among nuclei. Clusters were annotated based on the expression of established canonical marker genes for major cell types. Cell type composition was quantified as the proportion of nuclei assigned to each annotated cluster per sample.

### Crosslinking of W6/32 clone MHC Class I antibody to protein A-sepharose 4B beads

5 mg of antibody was loaded on 1 mL of Protein A-Sepharose 4B beads (Invitrogen) packed in a polypropylene column (BioRad) and incubated for 30 min at room temperature. Antibody-bound beads were washed with borate buffer (pH 9) and incubated with 20 mM dimethyl pimelimidate (DMP) linker for 45 min. Crosslinking was halted by incubating the column with ethanolamine for 2 hours. The column was washed with PBS and stored in PBS containing 0.02% sodium azide at 4°C. Prior to use, the column was washed with 0.1 N acetic acid and equilibrated with 100 mM Tris-HCl, pH 8.0.

### MHC-peptide complex enrichment

Human neural aggregate organoids were treated with IFNγ (100 ng/mL) for 72 hours. In some cases, 7-10 days prior to IFNγ treatment mature the organoids were transduced with adeno associated viral vectors (AAV1-SYN1.hB2M/T2A/EGFP:oPRE, VectorBuilder; AAV1.Syn.eGFP:WPRE, AddGene 50465-AAV1; AAV1.Syn.sig-CEF-DC-LAMP-T2A-eGFP.WPRE, Vector Biolabs). To obtain sufficient protein lysates for immunopeptidomic analysis 120-200 million neural cells were used for each condition. After IFNγ treatment, cells were washed on ice with cold PBS. Flasks were immediately scraped and cells were pelleted by centrifugation, snap frozen in liquid nitrogen, and stored until further processing. MHC-peptide complexes were enriched as described [25].

Human neural aggregate organoids were lysed in buffer containing 0.25% sodium deoxycholate, 0.2 mM IAA, 1 mM EDTA, 1 mM PMSF, 1% octyl-B-glucopyranoside, and 1:200 protease inhibitor cocktail for 1 h on ice. Cell debris was removed by centrifugation at 25,000 x g at 4°C for 50 min. The lysate was loaded onto affinity columns containing W6/32 crosslinked to protein A beads and incubated for 1 hour with gentle rotation. The column was washed sequentially with 1) 150 mM NaCl in 20 mM Tris-HCl pH 8.0, 2) 400 mM NaCl in 20 mM Tris-HCl pH 8.0, 3) 150 mM NaCl in 20 mM Tris-HCl pH 8.0, and 4) 20 mM Tris-HCl pH 8.0. MHC-bound peptide complexes were eluted using 1% TFA and the eluate was loaded on Sep-Pak C18 columns to purify the eluted peptides. The peptides were dried using a Speedvac concentrator prior to LC-MS/MS analysis.

### LC-MS/MS analysis

LC-MS/MS analysis was carried out on an Orbitrap Eclipse Tribrid mass spectrometer (Thermo Scientific) connected online to a Dionex RSLC3000 liquid chromatography system (Thermo Scientific). Peptides were suspended in solvent A (0.1% formic acid) and loaded on a trap column (PepMap C_18_ 2 cm × 100 µm, 100 Å) followed by high resolution separation column (EasySpray 50 cm X 75 µm, C_18_ 1.9 µm, 100 Å, Thermo Scientific). The mass spectrometer was operated in data dependent mode with a cycle time of 2 s. Survey MS scan was acquired in Orbitrap mass analyzer with 120K resolution, 4xe5 AGC target and 50 ms injection time. Monoisotopic precursor ions with charge state 2-4 were subjected to MS/MS with top scan priority followed by precursor with charge 1. Precursor ions (z=2-4) were fragmented with 28% HCD normalized collision energy and acquired in orbitrap mass analyzer with 15K resolution. Precursor ions (z = 1) with a mass range of 700 – 1400 m/z were fragmented with 32% HCD normalized collision energy and analyzed in orbitrap analyzer. Dynamic exclusion was enabled with 30 s exclusion duration. Additional filters included monoisotopic precursor selection and intensity threshold of 2.5×10^4^.

### Immunofluorescence and confocal microscopy

Neural aggregate organoids were initially fixed by adding an equal volume of 4% paraformaldehyde (PFA) prepared in 0.1 M phosphate buffer (pH 7.4) directly to the culture medium and incubating for 25 minutes at room temperature. The medium was aspirated and replaced with ice-cold 4% PFA in phosphate buffer for another 25 minutes. Cells were permeabilized in 0.1% Triton X-100 in PBS and blocked for at least 30 minutes in PBS containing 5% normal donkey serum, 1% bovine serum albumin, 0.01% sodium azide, and 0.1% Triton X-100. Cells were subsequently incubated overnight at 4°C with primary antibodies (anti-HLA-ABC (W6/32; Thermo Fisher), anti-GFAP (2.2B10; Thermo Fisher), anti-MAP2 (AP20; Millipore Sigma), anti-neuron-specific beta-III tubulin (MAB1195; R&D Systems)) diluted to 5 µg/mL in blocking buffer. The following day, cells were washed three times with 0.1% Triton X-100 in PBS and incubated for 2 hours at room temperature with secondary antibodies diluted 1:200 in blocking buffer. After washing, nuclei were counterstained with DAPI prior to imaging. Images were acquired using an Olympus confocal microscope and post-processed in ImageJ and Photoshop.

### Western blot

Protein samples were prepared in urea-based sample buffer (8 M urea, 50 mM Tris-HCl pH 7.5, 75 mM NaCl) supplemented with protease and phosphatase inhibitors. Samples were mixed with SDS-containing loading buffer supplemented with 50 mM DTT and resolved by SDS-PAGE on 4-12% gradient gels. Proteins were transferred onto 0.2 µm nitrocellulose membranes using wet transfer at 100 V for 90 min. Membranes were briefly stained with Ponceau S to confirm equal transfer and loading, then blocked in 5% BSA in TBS-T (Tris-buffered saline, 0.1% Tween-20) for 1 h at room temperature. Membranes were incubated overnight at 4°C with primary antibodies (anti-HLA-ABC (W6/32; Thermo Fisher), anti-HLA-A (EP1395Y; Abcam), anti-HLA-B (23GB6375; Thermo Fisher), anti-HLA-C (EPR6749; Abcam)) diluted in blocking buffer, washed three times in TBS-T, and incubated for 1 h at room temperature with HRP-conjugated secondary antibodies.

Immunoreactive bands were detected using enhanced chemiluminescence (ECL) substrate and imaged with a ChemiDoc imaging system (Bio-Rad). Band intensities were quantified using Image Lab software (Bio-Rad).

## DECLARATIONS

### Ethics approval and consent to participate

All human samples were obtained from individuals providing written informed consent following protocols approved by the Mayo Clinic Institutional Review Board (IRB 17-004547; IRB 13-007298).

### Consent for publication

Not applicable.

### Availability of data and materials

The datasets generated and/or analysed during the current study are available from the corresponding author on reasonable request. Data are not yet publicly available because they are being prepared for deposition in appropriate public repositories; accession numbers/DOIs will be provided upon publication. Requests that involve potentially identifiable donor-level information will require an appropriate data use agreement and applicable approvals.

### Author contributions

BDSC and CLH conceived and designed the study. BDSC, CLH, AP, KM, MC, and RR developed the methodology; BDSC, KM, RBS, SP, KM, and RR carried out the investigation and performed initial analysis of the data; BDSC generated initial figure drafts. BDSC and CLH validated the analyses and generated the final figures. CLH acquired funding, provided project administration, and supervised the project. BDSC, SP, and CLH wrote, edited, and approved the manuscript. All authors approved the final manuscript.

## Acknowledgements

We acknowledge the assistance of the Center for Multiple Sclerosis and Autoimmune Neurology at Mayo Clinic for providing patient PBMCs and fibroblasts and we thank the coordinators from the center for their efforts. We also acknowledge assistance of the Mayo Clinic Proteomics Core, which is a shared resource of the Mayo Clinic Cancer Center (NCI P30 CA15083). Dr. Parijat Kabiraj provided experimental support during early stages of the project. Brittany Overlee, Jonghoon Choi, and Henry Nguyen provided exceptional technical support.

## REFERENCES

1. Segal Y, Soltys J, Clarkson BDS, Howe CL, Irani SR, Pittock SJ: Toward curing neurological autoimmune disorders: Biomarkers, immunological mechanisms, and therapeutic targets. Neuron 2025, 113:345–379.

2. Eshaghi A, Prados F, Brownlee WJ, Altmann DR, Tur C, Cardoso MJ, De Angelis F, van de Pavert SH, Cawley N, De Stefano N, et al: Deep gray matter volume loss drives disability worsening in multiple sclerosis. Ann Neurol 2018, 83:210–222.

3. Schlaeger R, Papinutto N, Panara V, Bevan C, Lobach IV, Bucci M, Caverzasi E, Gelfand JM, Green AJ, Jordan KM, et al: Spinal cord gray matter atrophy correlates with multiple sclerosis disability. Ann Neurol 2014, 76:568–580.

4. Babbe H, Roers A, Waisman A, Lassmann H, Goebels N, Hohlfeld R, Friese M, Schroder R, Deckert M, Schmidt S, et al: Clonal expansions of CD8(+) T cells dominate the T cell infiltrate in active multiple sclerosis lesions as shown by micromanipulation and single cell polymerase chain reaction. J Exp Med 2000, 192:393–404.

5. Skulina C, Schmidt S, Dornmair K, Babbe H, Roers A, Rajewsky K, Wekerle H, Hohlfeld R, Goebels N: Multiple sclerosis: brain-infiltrating CD8+ T cells persist as clonal expansions in the cerebrospinal fluid and blood. Proc Natl Acad Sci U S A 2004, 101:2428–2433.

6. Neumann H, Medana IM, Bauer J, Lassmann H: Cytotoxic T lymphocytes in autoimmune and degenerative CNS diseases. Trends Neurosci 2002, 25:313–319.

7. Frischer JM, Weigand SD, Guo Y, Kale N, Parisi JE, Pirko I, Mandrekar J, Bramow S, Metz I, Bruck W, et al: Clinical and pathological insights into the dynamic nature of the white matter multiple sclerosis plaque. Ann Neurol 2015, 78:710–721.

8. Klistorner S, Barnett MH, Yiannikas C, Barton J, Parratt J, You Y, Graham SL, Klistorner A: Expansion of chronic lesions is linked to disease progression in relapsing-remitting multiple sclerosis patients. Mult Scler 2021, 27:1533–1542.

9. Terrabuio E, Pietronigro EC, Bani A, Della Bianca V, Laudanna C, Rossi B, Finotti G, Santos-Lima B, Zenaro E, Turano E, et al: CD103(-)CD8(+) T cells promote neurotoxic inflammation in Alzheimer’s disease via granzyme K-PAR-1 signaling. Nat Commun 2025, 16:8372.

10. Kim HJ, Ban JJ, Kang J, Im HR, Ko SH, Sung JJ, Park SH, Park JE, Choi SJ: Single-cell analysis reveals expanded CD8(+) GZMK (high) T cells in CSF and shared peripheral clones in sporadic amyotrophic lateral sclerosis. Brain Commun 2024, 6:fcae428.

11. Su W, Saravia J, Risch I, Rankin S, Guy C, Chapman NM, Shi H, Sun Y, Kc A, Li W, et al: CXCR6 orchestrates brain CD8(+) T cell residency and limits mouse Alzheimer’s disease pathology. Nat Immunol 2023, 24:1735–1747.

12. Chen X, Firulyova M, Manis M, Herz J, Smirnov I, Aladyeva E, Wang C, Bao X, Finn MB, Hu H, et al: Microglia-mediated T cell infiltration drives neurodegeneration in tauopathy. Nature 2023, 615:668–677.

13. Campisi L, Chizari S, Ho JSY, Gromova A, Arnold FJ, Mosca L, Mei X, Fstkchyan Y, Torre D, Beharry C, et al: Clonally expanded CD8 T cells characterize amyotrophic lateral sclerosis-4. Nature 2022, 606:945–952.

14. Gate D, Saligrama N, Leventhal O, Yang AC, Unger MS, Middeldorp J, Chen K, Lehallier B, Channappa D, De Los Santos MB, et al: Clonally expanded CD8 T cells patrol the cerebrospinal fluid in Alzheimer’s disease. Nature 2020, 577:399–404.

15. Clarkson BDS, Patel MS, LaFrance-Corey RG, Howe CL: Retrograde interferon-gamma signaling induces major histocompatibility class I expression in human-induced pluripotent stem cell-derived neurons. Ann Clin Transl Neurol 2018, 5:172–185.

16. Clarkson BD, Grund EM, Standiford MM, Mirchia K, Westphal MS, Muschler LS, Howe CL: CD8+ T cells recognizing a neuron-restricted antigen injure axons in a model of multiple sclerosis. J Clin Invest 2023, 133.

17. Kubiniok P, Marcu A, Bichmann L, Kuchenbecker L, Schuster H, Hamelin DJ, Duquette JD, Kovalchik KA, Wessling L, Kohlbacher O, et al: Understanding the constitutive presentation of MHC class I immunopeptidomes in primary tissues. iScience 2022, 25:103768.

18. Girdlestone J, Isamat M, Gewert D, Milstein C: Transcriptional regulation of HLA-A and -B: differential binding of members of the Rel and IRF families of transcription factors. Proc Natl Acad Sci U S A 1993, 90:11568–11572.

19. Brynedal B, Duvefelt K, Jonasdottir G, Roos IM, Akesson E, Palmgren J, Hillert J: HLA-A confers an HLA-DRB1 independent influence on the risk of multiple sclerosis. PLoS One 2007, 2:e664.

20. Wissemann WT, Hill-Burns EM, Zabetian CP, Factor SA, Patsopoulos N, Hoglund B, Holcomb C, Donahue RJ, Thomson G, Erlich H, Payami H: Association of Parkinson disease with structural and regulatory variants in the HLA region. Am J Hum Genet 2013, 93:984–993.

21. Paul S, Weiskopf D, Angelo MA, Sidney J, Peters B, Sette A: HLA class I alleles are associated with peptide-binding repertoires of different size, affinity, and immunogenicity. J Immunol 2013, 191:5831–5839.

22. Schellens IM, Hoof I, Meiring HD, Spijkers SN, Poelen MC, van Gaans-van den Brink JA, van der Poel K, Costa AI, van Els CA, van Baarle D, Kesmir C: Comprehensive Analysis of the Naturally Processed Peptide Repertoire: Differences between HLA-A and B in the Immunopeptidome. PLoS One 2015, 10:e0136417.

23. Mangalaparthi KK, Madugundu AK, Ryan ZC, Garapati K, Peterson JA, Dey G, Prakash A, Pandey A: Digging deeper into the immunopeptidome: characterization of post-translationally modified peptides presented by MHC I. J Proteins Proteom 2021, 12:151–160.

24. Chong C, Coukos G, Bassani-Sternberg M: Identification of tumor antigens with immunopeptidomics. Nat Biotechnol 2022, 40:175–188.

25. Bassani-Sternberg M, Braunlein E, Klar R, Engleitner T, Sinitcyn P, Audehm S, Straub M, Weber J, Slotta-Huspenina J, Specht K, et al: Direct identification of clinically relevant neoepitopes presented on native human melanoma tissue by mass spectrometry. Nat Commun 2016, 7:13404.

26. Nilsson JB, Greenbaum J, Peters B, Nielsen M: NetMHCpan-4.2: improved prediction of CD8+ epitopes by use of transfer learning and structural features. Front Immunol 2025, 16:1616113.

27. Reynisson B, Alvarez B, Paul S, Peters B, Nielsen M: NetMHCpan-4.1 and NetMHCIIpan-4.0: improved predictions of MHC antigen presentation by concurrent motif deconvolution and integration of MS MHC eluted ligand data. Nucleic Acids Res 2020, 48:W449–W454.

28. Nulman J, Ulrich JD, Holtzman DM: Context matters: Conflicting roles of interferon-gamma signaling in CNS diseases. Curr Opin Neurobiol 2025, 96:103149.

29. Carre A, Samassa F, Zhou Z, Perez-Hernandez J, Lekka C, Manganaro A, Oshima M, Liao H, Parker R, Nicastri A, et al: Interferon-alpha promotes HLA-B-restricted presentation of conventional and alternative antigens in human pancreatic beta-cells. Nat Commun 2025, 16:765.

30. Rizvi SM, Salam N, Geng J, Qi Y, Bream JH, Duggal P, Hussain SK, Martinson J, Wolinsky SM, Carrington M, Raghavan M: Distinct assembly profiles of HLA-B molecules. J Immunol 2014, 192:4967–4976.

31. Geng J, Zaitouna AJ, Raghavan M: Selected HLA-B allotypes are resistant to inhibition or deficiency of the transporter associated with antigen processing (TAP). PLoS Pathog 2018, 14:e1007171.

32. Loonstra FC, de Ruiter LRJ, Koel-Simmelink MJA, Schoonheim MM, Strijbis EMM, Moraal B, Barkhof F, Uitdehaag BMJ, Teunissen C, Killestein J: Neuroaxonal and Glial Markers in Patients of the Same Age With Multiple Sclerosis. Neurol Neuroimmunol Neuroinflamm 2023, 10.

33. Bittner S, Oh J, Havrdova EK, Tintore M, Zipp F: The potential of serum neurofilament as biomarker for multiple sclerosis. Brain 2021, 144:2954–2963.

34. Disanto G, Barro C, Benkert P, Naegelin Y, Schadelin S, Giardiello A, Zecca C, Blennow K, Zetterberg H, Leppert D, et al: Serum Neurofilament light: A biomarker of neuronal damage in multiple sclerosis. Ann Neurol 2017, 81:857–870.

35. Verde F, Otto M, Silani V: Neurofilament Light Chain as Biomarker for Amyotrophic Lateral Sclerosis and Frontotemporal Dementia. Front Neurosci 2021, 15:679199.

36. Puentes F, Topping J, Kuhle J, van der Star BJ, Douiri A, Giovannoni G, Baker D, Amor S, Malaspina A: Immune reactivity to neurofilament proteins in the clinical staging of amyotrophic lateral sclerosis. J Neurol Neurosurg Psychiatry 2014, 85:274–278.

37. Puentes F, van der Star BJ, Boomkamp SD, Kipp M, Boon L, Bosca I, Raffel J, Gnanapavan S, van der Valk P, Stephenson J, et al: Neurofilament light as an immune target for pathogenic antibodies. Immunology 2017, 152:580–588.

38. Huizinga R, Hintzen RQ, Assink K, van Meurs M, Amor S: T-cell responses to neurofilament light protein are part of the normal immune repertoire. Int Immunol 2009, 21:433–441.

39. Huizinga R, Gerritsen W, Heijmans N, Amor S: Axonal loss and gray matter pathology as a direct result of autoimmunity to neurofilaments. Neurobiol Dis 2008, 32:461–470.

40. Clarkson BD, Johnson RK, Bingel C, Lothaller C, Howe CL: Preservation of antigen-specific responses in cryopreserved CD4(+) and CD8(+) T cells expanded with IL-2 and IL-7. J Transl Autoimmun 2022, 5:100173.

